# Missense Mutant Gain-of-Function Causes Inverted Formin 2 (INF2)-Related Focal Segmental Glomerulosclerosis (FSGS)

**DOI:** 10.1101/2024.06.08.598088

**Authors:** Balajikarthick Subramanian, Sarah Williams, Sophie Karp, Marie-Flore Hennino, Sonako Jacas, Miriam Lee, Cristian V. Riella, Seth L. Alper, Henry N. Higgs, Martin R. Pollak

## Abstract

Inverted formin-2 (INF2) gene mutations are among the most common causes of genetic focal segmental glomerulosclerosis (FSGS) with or without Charcot-Marie-Tooth (CMT) disease. Recent studies suggest that INF2, through its effects on actin and microtubule arrangement, can regulate processes including vesicle trafficking, cell adhesion, mitochondrial calcium uptake, mitochondrial fission, and T-cell polarization. Despite roles for INF2 in multiple cellular processes, neither the human pathogenic R218Q INF2 point mutation nor the INF2 knock-out allele is sufficient to cause disease in mice. This discrepancy challenges our efforts to explain the disease mechanism, as the link between INF2-related processes, podocyte structure, disease inheritance pattern, and their clinical presentation remains enigmatic. Here, we compared the kidney responses to puromycin aminonucleoside (PAN) induced injury between R218Q INF2 point mutant knock-in and INF2 knock-out mouse models and show that R218Q INF2 mice are susceptible to developing proteinuria and FSGS. This contrasts with INF2 knock-out mice, which show only a minimal kidney phenotype. Co-localization and co-immunoprecipitation analysis of wild-type and mutant INF2 coupled with measurements of cellular actin content revealed that the R218Q INF2 point mutation confers a gain-of-function effect by altering the actin cytoskeleton, facilitated in part by alterations in INF2 localization. Differential analysis of RNA expression in PAN-stressed heterozygous R218Q INF2 point-mutant and heterozygous INF2 knock-out mouse glomeruli showed that the adhesion and mitochondria-related pathways were significantly enriched in the disease condition. Mouse podocytes with R218Q INF2, and an INF2-mutant human patient’s kidney organoid-derived podocytes with an S186P INF2 mutation, recapitulate the defective adhesion and mitochondria phenotypes. These results link INF2-regulated cellular processes to the onset and progression of glomerular disease. Thus, our data demonstrate that gain-of-function mechanisms drive INF2-related FSGS and explain the autosomal dominant inheritance pattern of this disease.

## INTRODUCTION

Glomerular epithelial cells, or podocytes, are highly specialized cells that wrap around the outer surface of glomerular capillaries ^1, 2^. They exhibit a unique morphology, wherein the cell body gives rise to long extensions that branch into primary processes and secondary foot processes. These foot processes interdigitate with the foot processes of neighboring podocytes to form filtration slit diaphragms. This complex morphology depends on the underlying actin cytoskeletal arrangement, which, if impaired, can lead to podocyte injury and kidney disease ^3, 4^.

Focal segmental glomerulosclerosis (FSGS) is a histological pattern of injury defined by the presence of scarring (sclerosis) in some parts (segmental) of certain glomeruli (focal) within the kidney ^5^. This pathology occurs when podocytes are dedifferentiated or reduced in number due to injury ^5, 6^. To date, mutations in more than 50 genes have been described as the causative events affecting the podocytes’ structure directly or indirectly ^7, 8^. Of these, mutations in an actin regulatory gene, Inverted Formin 2 (INF2), are relatively common, accounting for 17% of familial FSGS and 1% of sporadic cases ^9, 10, 11, 12^. Over 60 different disease-associated INF2 mutations have been identified ^12^. These mutations lead to a kidney phenotype characterized by proteinuria, progressive kidney dysfunction, and FSGS with or without Charcot–Marie Tooth disease (CMT) ^11, 13^.

INF2 is one of the 15-member formin family of proteins that are primarily involved in polymerizing monomeric ‘globular’ actin (G-actin) into actin filaments (F-actin) ^14, 15^. The defining feature of these formins, including INF2, is a formin homology domain (FH2) responsible for actin nucleation and elongation. In addition, they contain other domains, such as formin homology 1 (FH1) and diaphanous autoregulatory domain (DAD), working in tandem with FH2 to promote actin assembly. Importantly, the amino-terminal region of INF2 contains a diaphanous inhibitory domain (DID) that can interact with the DAD in the C-terminal region of INF2 and enable the protein to fold to an autoinhibited state ^12, 14^ ^16^ which tightly regulates actin assembly. All identified FSGS-causing mutations in INF2 localize to within INF2-DID, leading to the release of the DAD and other domains from this inhibition ^12^.

Over the past several years, multiple studies have demonstrated roles for INF2 in mitochondrial calcium up-take, mitochondrial fission, cell adhesion, vesicle trafficking, T-cell polarization, placental implantation, and numerous other cell processes ^12, 17, 18, 19, 20, 21^ ^22^. Although these studies highlight INF2’s importance in cells, gaps persist in understanding its role in regulating podocytes’ unique structure and function, limiting our understanding of the related FSGS disease mechanism.

In this study, to better define the mechanism(s) of INF2-related kidney disease, we interrogated R218Q INF2 point mutant (knock-in) and INF2 knock-out mouse models in response to puromycin aminonucleoside (PAN) stress and compared their phenotypic responses. We report that, in contrast to the INF2 knock-out mouse model, the R218Q knock-in mutant mouse model is susceptible to developing proteinuria and FSGS in response to PAN injury. Furthermore, the ability of R218Q INF2 to alter the localization of wild-type INF2 and induce defects in actin arrangements, podocyte adhesion, and mitochondria indicates that a gain-of-function mechanism drives the development of disease. These data help explain the autosomal dominant inheritance of this disease.

## Materials and Methods

### Ethical Statement

All practices involving mice were performed in compliance with an animal care protocol approved by the Institute Animal Care and Use Committee (IACUC) at Beth Israel Deaconess Medical Center.

### Mouse Experiments

PAN injury response in INF2 mouse models was assessed as described previously ^23^. Briefly, the mice were injected with PAN intraperitoneally (490 mg/kg), and urine samples were collected on Day 0 (before the PAN injection), 3, 7, 11, and 14 for analysis. The SDS-PAGE gel electrophoresis method was used to separate the urine samples, which were then stained with Coomassie dye to detect albuminuria. Albumin and creatinine levels in the urine were quantified using an Albumin ELISA Kit (Bethyl Labs, TX) and a colorimetric kit, respectively, following the manufacturers’ protocols.

### Histology

Mice kidneys were harvested at indicated time frames and processed to examine glomerulosclerosis and podocyte foot process arrangement. For analyzing sclerosis, kidneys were fixed in 10% buffered formalin and paraffin-embedded to prepare FFPE blocks. These blocks were then cut into five-micron sections and processed for periodic acid-Schiff and Masson’s trichrome staining. The ratio of sclerotic glomeruli to total glomeruli was calculated and plotted. For podocyte ultrastructure analysis, kidneys were fixed in a modified Karnovsky’s fixative (2.5% glutaraldehyde and 2% paraformaldehyde in 0.1M Cacodylate buffer, pH 7.4), processed using 1% osmium tetroxide and Epon-embedded for sectioning. Sections collected on grids were imaged by the JEOL 1400 Transmission electron microscope. The number of foot processes per micron was quantified and plotted.

### Cell Culture

Mouse podocytes, derived from INF2 mouse models (wild-type, heterozygous R218Q knock-in, heterozygous knock-out, homozygous R218Q knock-in, homozygous knock-out), were maintained in RPMI 1640 (Corning, MA), supplemented with 1% Insulin-Transferrin-Selenium (ITS), 1% penicillin-streptomycin, and 10% Fetal Bovine Serum (FBS) ^24^. Human embryonic kidney epithelial cells (HEK293T cells) were maintained in DMEM (Corning, MA) supplemented with 10% FBS and 1% penicillin-streptomycin.

### INF2 Plasmid Constructs and Expression

For the RFP-tagged construct, the full-length human INF2-CAAX isoform was cloned into a Turbo RFP plasmid (N-terminal RFP) (Origene, MD). For GFP-tagged constructs, human INF2-CAAX isoform sequences corresponding to amino acids 1–547 (N-fragment), 1–1249 (full-length) were cloned into an EGFPC1 plasmid (N-terminal GFP tag) (Takara Bio, CA). For non-cleavable INF2, amino acids 543–548 of INF2 were replaced with hexa-alanine in full-length CAAX constructs. For HA and Flag-tagged construct, full-length INF2-CAAX isoform was cloned into pCMV6 vector. HA-tag was included at the N-terminus, and Flag-tag was added at the C-terminus of INF2. For transient expression, plasmids were transfected with lipofectamine 3000 (for podocytes) or 2000 (for 293T cells) as per the manufacturer’s guidelines (Thermo Fisher Scientific, MA). All interaction and imaging analyses were performed at 48 hours post-transfection. For INF2 constructs for the pyrene-actin assays, human INF2-FL-nonCAAX cDNA was cloned into an EGFPC1 vector containing Strep-tag II (IBA Life Sciences) and an HRV3C cleavage site that was located N-terminal to the INF2 start codon. For INF2 DID proteins (wild-type and R218Q), human INF2 N-terminus region (amino acids 1-267) was cloned into pGEX-KT vector.

### Coimmunoprecipitation

HEK293T cells were transiently transfected with the indicated INF2 constructs using Lipofectamine 2000 (Invitrogen, CA). After 48 hours, the cells were lysed in 1% NP-40 lysis buffer (1% NP-40, 50 mM Tris-HCl, 150 mM NaCl, 5 mM EDTA, pH 7.4) supplemented with protease inhibitors. Cell lysates were then incubated with mouse anti–FLAG beads (MA5–15256; ThermoFisher, MA) for 4 hours, followed by their magnetic sorting and suspension in premixed Laemmli buffer (BioRad, CA). The coimmunoprecipitation of full-length HA-INF2-FLAG with various GFP-tagged INF2 forms was then analyzed. Samples were probed for HA or GFP using the antibodies for GFP (1:500) (ab13970; Abcam, MA) or HA (1:500) (2367; Cell Signaling Technology, MA). The membranes were then incubated with IRDye secondary antibodies and imaged using a Odyseey Clx Infrared system (LI-COR, NE).

### Immunoblotting

Cell lysates were prepared in a RIPA buffer (Boston BioProducts, MA) supplemented with a cocktail of protease and phosphatase inhibitors (Roche, CA). Equal protein loads were electrophoresed on a 4%–20% gradient gel, quick-transferred to a polyvinylidene difluoride membrane (Bio-Rad, CA), and probed with respective primary and secondary antibodies as follows: INF2 (1:500) (A303–427A; Bethyl Laboratories, TX); INF2 (C) (1:500) (20466–1-AP; Proteintech, IL); Podocin (1:1000) (P0372; Sigma, MO); and anti-rabbit (1:4000) (Santa Cruz Biotechnology, CA). The membranes were then developed using a chemiluminescent-based substrate (Super Signal West Dura). Total *β*-actin (1:4000) (sc47778; Santa Cruz Biotechnology, CA) level was used as a loading control.

### Immunofluorescence

Cells were PBS-rinsed and fixed with 4% para-formaldehyde for 15 minutes. The fixed cells were then quenched, permeabilized with 0.5% triton x-100, and blocked using 5% BSA solution. The primary and secondary antibody preparations were sequentially incubated for 2hr and counterstained cell nuclei with Hoechst (dsDNA) (Invitrogen). For organoid and mouse glomeruli, samples were processed into paraffin blocks, and five-micron sections were then cut and processed for staining analysis. The following primary antibodies with indicated dilutions were prepared in blocking solution and used for staining analysis: Nephrin (BP5030, 1:200; Origene, MD), INF2 (DT-157, 1:100; ref[ ]), WT-1 (ab89901, 1:100; Abcam, MA), a-SMA (ab56941, 100; Abcam, MA), paxillin (ab32084, 1:100; Abcam, MA), synaptopodin (AP33487SU-N, 1:200; Origene, MD), cortactin (MA5-15831, 1:100; Thermo Fisher Scientific, MA) F-actin (Rhodamine-Phalloidin, 1:100; Invitrogen, CA). For mitochondrial imaging, cells were incubated with deep red Mito tracker dye (8778; Cell Signaling Technology, MA) per the manufacturer’s instructions and imaged as live cells. All samples were imaged for fluorescence using laser-scanning confocal microscopy (LSM 510; Zeiss) using ZEN lite Black edition software.

### Micropatterning cells

Cross-bow micropatterns, custom prepared in Chromemask (Advance Repro, MA), were used to program cells to reach uniform shape and size. Micropatterning of cells using the mask was performed as described previously ^25^. Briefly, poly-l-lysine-grafted-polyethylene glycol (PLL-g-PEG) was coated on a cover glass, placed under a chrome mask containing the micropatterns, and exposed to a UV light through the chrome mask. The exposed parts of the PLL-PEG were washed in distilled water, dried, and coated with fibronectin (250 ng/mL). Cells were plated as single suspensions. After 1 hr, cover glasses were washed away to remove unattached cells. For PAN treatments, cells were pre-treated with PAN overnight, trypsinized and replated with PAN on micropatterns and maintained through cell attachment. All analyses were performed after six hours of cell spreading in micropatterns.

### Actin Assays

Mouse podocytes were lysed and examined for their total G-actin and F-actin levels using a G-actin/F-actin In vivo Biochemical Assay Kit (Cytoskeleton, CO). Briefly, 100 µl of precleared lysates were ultracentrifuged for 100,000 g at 37C for 1hr. The supernatant containing the G-actin was collected and processed for immunoblotting. The residual pellet containing the F-actin was incubated with 100 µl of depolymerizing buffer in ice for 1 hr and then processed further for immunoblotting. The ratio of F-actin to G-actin was calculated from their band intensity.

### Biochemical actin polymerization assays

Actin polymerizing activity of full-length INF2 with wild-type and R218Q mutant DID regions was assessed using pyrene actin polymerization assay as described previously ^26^. Briefly, rabbit skeletal muscle actin in G-buffer (6 μM actin, 10% pyrene) were converted to Mg2+ salt by the addition of EGTA and MgCl2 (to 1 and 0.1 mM, respectively) for 2 min at 23°C immediately before polymerization. Polymerization was induced by two volumes of 1.5xpolymerization buffer (75mM KCl, 1.5mM MgCl2, 1.5mM EGTA, 15 mM Hepes pH 7.4, 2mM DTT, 2 mM Tris-HCl, 0.2 mM ATP, 0.1 mMCaCl2, and 0.01% w/v NaN3) containing other proteins (20 nM INF2-FL, 6 μM profilin, 50 nM capping protein and/or 0.5 μM DID region). Pyrene fluorescence (365/410 nm) was monitored in a 96-well fluorescence plate reader within 1 minute of inducing polymerization and plotted versus time.

### RNA-seq and Gene Set Enrichment Analysis

Total RNA from mouse glomeruli was sequenced on a HiSeq 4000 following standard workflow using NEBNext Ultra II RNA Library Preparation kit (New Englab Biolabs, MA)(Illumina, CA) and paired-end sequencing (Genewiz, MA). The samples had 78 million pass filter reads, with 89% of bases above the quality score of Q30 and a mean quality score of 37.32. Adapters and low-quality bases were trimmed and aligned with the mouse reference genome (GRCm38 ) and GENCODE annotation using STAR 2.7. The average mapping rate of all samples was 90%.

Genes showing differential expression were analyzed for (1) HET. KI versus HET. KO, (2) PAN-treated HET.KI versus PAN-treated HET.KO, (3) HET.KI versus PAN-treated HET.KI, and (4) HET.KO versus PAN-treated HET.KO groups. The common and unique genes between these analyses were identified using DiVenn 1.2. Gene set enrichment analysis was conducted between PAN-treated HET. KI versus PAN-treated HET. KO (GSEA v 4.3.3 https://www.gsea-msigdb.org/gsea/license_terms_list.jsp). The enriched gene ontology Cellular Component gene sets (p<0.05) were identified and visualized as a network in Cytoscape using EnrichmentMap, AutoAnnotate, and clusterMaker2 application. The overlap of the number of genes between gene set enrichment analysis and Divenn analysis was plotted using Venny 2.1.

### Human Kidney Organoids and Outgrown Podocyte analysis

Human IPSC cell lines were generated from a normal and INF2-related FSGS patient using CytoTune-iPS 2.0 Sendai reprogramming kit (ThermoFisher, MA) at Harvard Stem Cell Institute IPS Core Facility. Kidney organoids were then prepared as described previously^27^, with minor modifications. In each step where APEL was called for, APELII supplemented with 5% protein-free hybridomia medium II (PFHMII) (ThermoFisher, MA) was used. In addition, the initial nine days of the protocol were carried out in 3D gel suspension to enhance homogeneity in differentiation.

Briefly, 6 μl of the IPSc cell suspension was plated in ∼175 μl collagen matrix in 3-D in (222μl Rat tail collagen (∼10mg/mL) or 667μl Rat tail collagen (3mg/mL), 227μl geltrex, 100μl 10x PBS, 4 μl 1N NaOH) in 24 well plates with 1 mL mTeSR for 48 hours. The media was then changed to 0.5 mL APELII medium supplemented with 5% PFHMII and 8μM CHIR99021 for 4 days with media changes every two days. On day 4, media was switched to 1 mL of APELII supplemented with 5% PFHMII and 200 ng ml−1 FGF9 and 1 μg ml−1 heparin until it has been 7 days since the cells were first introduced to CHIR99021. The collagen mix was then digested with ∼200μl Collagenase A (10mg/mL) for 30 minutes. The released spheroids were spun down into aggregates with 1 mL APEL2 with 5% PFHMII, and all further steps were performed in accordance with the Takasato et al. 2016 protocol. For PAN treatments, mature organoids (5+ days post aggregate formation) were given APELII media with 5% PFHMII and 1 mg/mL PAN dissolved in PBS. For harvesting podocytes, ten organoid cultures were pooled, and glomeruli extraction was performed using the sieve method, as described previously ^28^. The isolated glomerular structures were then maintained in a media composition of RPMI 1640 supplemented with 10% fetal bovine serum and 1% ITS and podocytes were outgrown on a Collagen I-coated surface. After one week of culturing, outgrown podocytes were used for cell adhesion and mitochondria assessments.

### Plotting and Statistical Analysis

All plotting and statistical analyses were performed using OriginPro software. All values are reported as mean±SD. Analyses were performed using one-way ANOVA between test groups. Tukey’s multiple comparison test was used to find group differences. Statistical significance was set at a minimal value of p<0.05. For evaluating the correlation between mice features and disease development, PAN-stressed mice were categorized based on sex (male, female), age (mature adult (>6 months), middle aged (>12 month)), INF2 genotypes (wild-type, heterozygous knock-in, heterozygous knock-out, homozygous knock-in, homozygous knock-out), and analyzed using the Pearson correlation coefficient method for their correlation with phenotype development (proteinuria). Features with r values>0.5 is selected as having a strong positive correlation with disease development.

## RESULTS

### INF2–mediated FSGS is a gain-of-function disease

Prior human genetic studies have described pathogenic mutations at multiple positions within the INF2 DID ^12^. All of these FSGS-causing INF2 mutations are found in the heterozygous state, consistent with the dominant inheritance of the disease, and all encode missense amino acid changes. None are predicted to cause total protein loss. Interestingly, mice with complete knock-out of INF2 are grossly normal and lack any apparent kidney disease phenotype ^24^. Based on these observations, we reasoned that the INF2-mediated FSGS disease likely occurs via pure gain-of-function effect of mutant INF2.

To examine this experimentally, we compared heterozygous INF2 R218Q knock-in (point mutant) mice and heterozygous INF2 knock-out mice. We stressed these mice with PAN and compared the development of kidney disease as a function of genotype. When we examined the development of albuminuria (measured as urinary albumin to creatinine ratio), we observed significant albuminuria at day 3 in the heterozygous knock-in mouse model. The proteinuria peaked at day 11 post-PAN injury, after which levels of urine albumin declined but did not return to the pre-inury baseline level. In contrast, the heterozygous knock-out mouse model did not develop any albuminuria within the indicated time frames post-PAN injury (Figure 1A-B).

**Figure 1.**
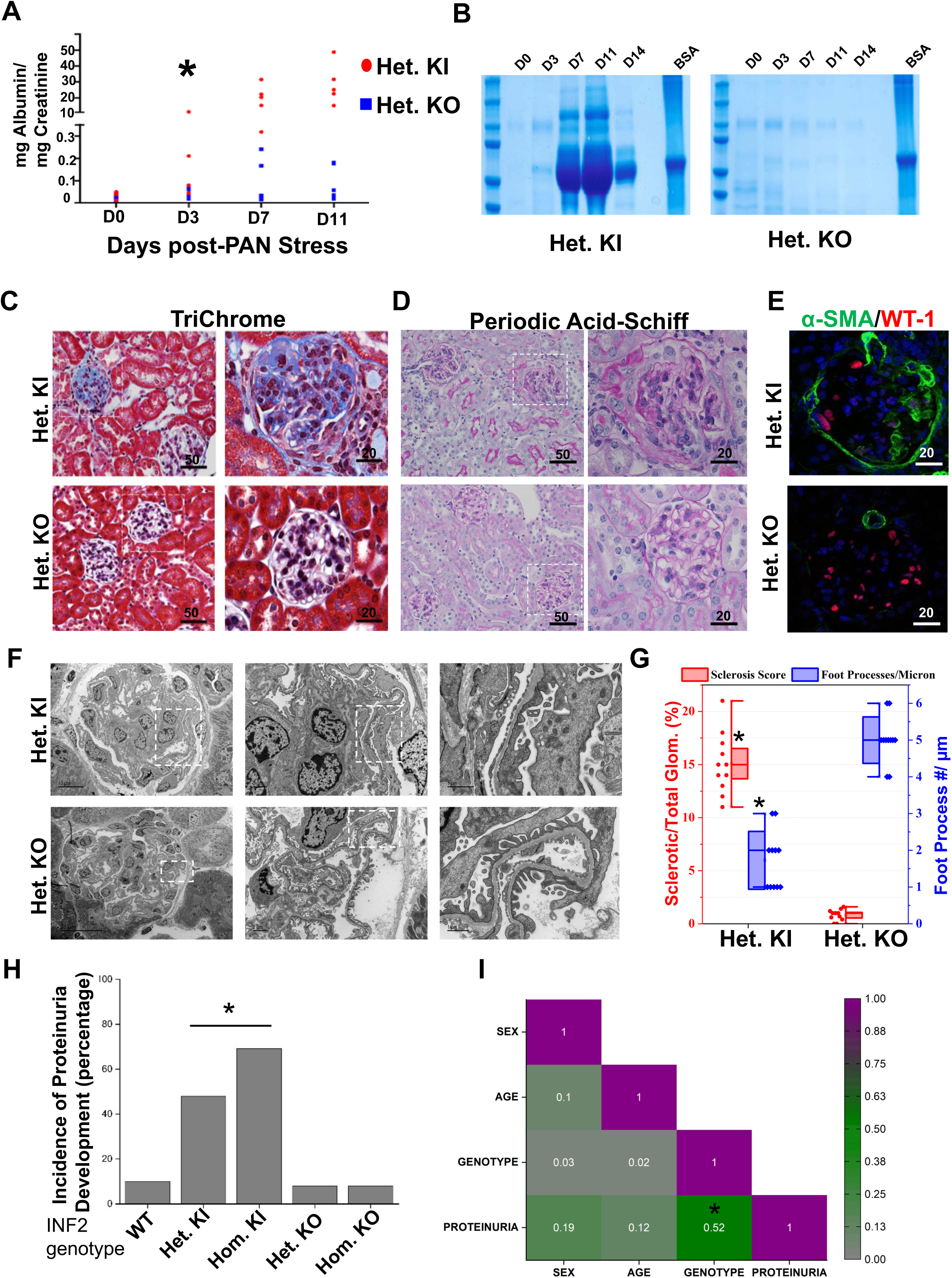
INF2 R218Q Knock-in mutant and INF2 Knock-out mouse models differ in disease development. (A-G) Heterozygous INF2 R218Q knock-in and heterozygous INF2 knock-out mouse models were stressed with puromycin aminonucleoside (PAN) and assessed for development of kidney disease. (A) Quantification of urinary albumin to creatinine ratio. Statistical significant differences were seen between knock-in versus knock-out mice from day 3. (*p<0.01; Student’s *t-*test) (B) Urine-gel electrophoresis. Urine samples from different time points of a representative PAN-stressed heterozygous knock-in and heterozygous knock-out mouse underwent electrophoresis in an SDS-PAGE gel. (C-D) Histology analysis. PAN-stressed mice were examined after eight weeks for histological changes. (C) Tri Chrome staining (D) Periodic Acid-Schiff staining. Fibrotic lesions were noted in heterozygous knock-in but not in heterozygous knock-out mice. Higher magnification images of the highlighted regions are shown in series. (E) Immunofluorescence analysis of WT-1(podocyte marker) and alpha-smooth muscle actin (fibrosis marker). Increased smooth muscle actin expression was noted in PAN-stressed heterozygous knock-in mice but not in heterozygous knock-out mice. (F) Ultrastructural analysis of podocytes. Higher magnification images of highlighted regions were shown in series. (G) Quantification of sclerosis and foot processes effacement. (*p<0.01; Student’s *t-*test). (H) Disease development among various INF2 genotypes. Mice from all INF2 genotypes were PAN-stressed and evaluated for proteinuria development. Incidence of proteinuria incidences were significantly higher in heterozygous and homozygous knock-in genotype mice (*p<0.01; one-way ANOVA). (I) Correlogram representing the Pearson correlation coefficient matrix between various mice features. Colors indicate the values of the correlation coefficient, as indicated in the scale bar. A significant correlation was present between genotype and proteinuria but not with other mice features (*p<0.05; r, 0.52). All scale bars are in microns.

We then assessed whether podocyte injury and related histological changes follow the pattern of albuminuria in the heterozygous knock-in and knock-out mouse models. We observed a fragmented pattern for marker proteins nephrin and endomucin, reduced WT-1 positive cells, and increased alpha-smooth muscle actin expression in PAN-injured heterozygous knock-in mouse glomeruli. These alterations were not present in the glomeruli of the heterozygous knock-out mouse model (Figure 1C and Supplementary Figure 1). To examine the disruption in the foot process/slit diaphragm and transition towards fibrosis, we assessed the kidney sections using Periodic Acid-Schiff (PAS) and Masson’s trichrome staining and electron microscopy. We observed fibrotic lesions and foot process effacement in PAN-stressed heterozygous R218Q knock-in mice, characteristic of an FSGS pattern of injury (Figure 1D-G). In contrast, the PAN stressed-heterozygous INF2 knock-out mouse did not develop these aberrations in marker protein distribution and glomerular histology (Supplementary Figure 1 and Figure 1C-G).

Next, we tested whether disease development in heterozygous knock-in mice was driven solely by the mutant allele’s gain-of-function effects. For this purpose, we subjected PAN stress to mice spanning a range of INF2 genotypes – wild-type, heterozygous, and homozygous forms of knock-in and knock-out alleles, genders, and adult age groups. We observed a high incidence of proteinuria in both heterozygous and homozygous knock-in mice, whereas both the heterozygous and homozygous knock-out mice had infrequent proteinuria, similar to wild-type mice (Figure 1H). Pearson correlation coefficients revealed a positive correlation between disease development and mutant allele genotype but not with other gender- and age-related features of the mice (Figure 1I). These observations indicate that INF2-mediated kidney disease occurs as a consequence of the gain-of-function effects of mutant INF2.

### Mutation in INF2-DID confers gain-of-function effect by altering the localization of wild-type INF2 and INF2 activity

Since INF2 is a member of the formin family of proteins that are primarily involved in regulating actin and the actin cytoskeleton, we reasoned that a DID-mutation-induced gain-of-function would likely manifest in alterations to the actin arrangement in cells. To assess this, we compared the F-actin/G-actin ratio in cells derived from mice of various genotypes in both untreated and PAN-treated podocytes. We observed that the knock-in mouse podocytes, particularly in homozygous conditions, exhibited a higher F-actin/G-actin ratio compared to wild-type and knock-out podocytes (Figure 2A). This trend is further amplified PAN treatment, causing a significant change in both heterozygous and homozygous knock-in podocytes.

**Figure 2.**
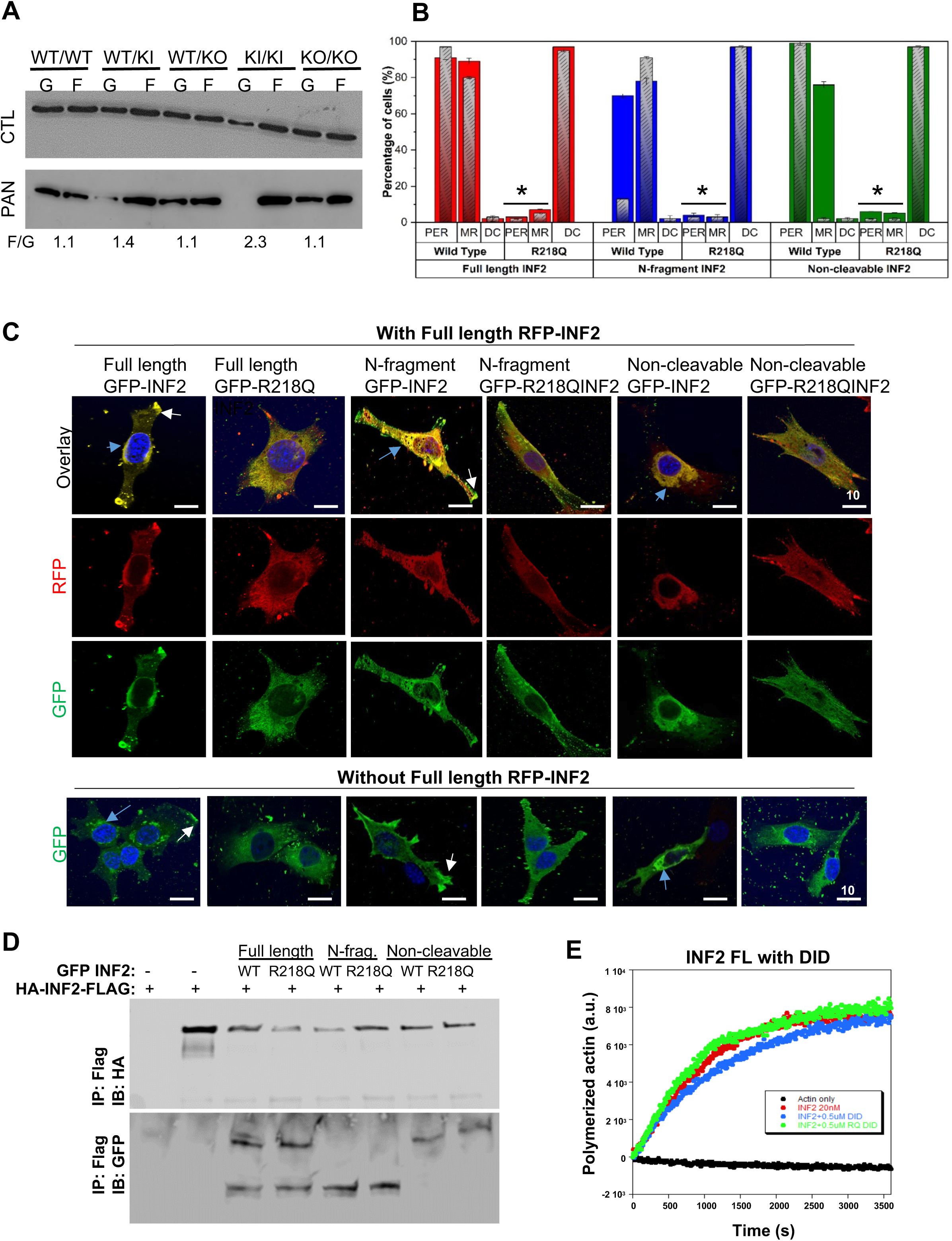
R218Q pathogenic mutation leads to altered localization and activity of wild-type INF2. (A) F/G actin assay. Immunoblot of β-actin. Control and PAN-stressed podocytes were examined for F-actin and G-actin content. Homozygous R218Q INF2 knock-in podocytes showed mild alteration in the basal condition in F-actin/G-actin distribution. In PAN-stressed conditions, both heterozygous and homozygous R218Q INF2 knock-in podocytes showed a prominent F-actin/G-actin distribution alteration. A representative immunoblot of three independent experiments was shown. The numerical value represents the mean F/G actin ratio for the PAN-stressed condition from three independent experiments. WT, wild-type; KI, knock-in; KO, knock-out. (B-C) Wild-type and R218Q GFP-tagged INF2 forms were co-expressed with or without RFP-tagged wild-type full-length INF2. Wild-type full-length RFP-INF2 showed a peri-nuclear ER-like pattern (blue arrow) with some membrane regions (white arrow highlights – membrane region localization). The coexistence of R218Q GFP INF2 forms (full-length, N-fragment, non-cleavable) alters this localization to a diffused cytoplasmic pattern. (B) Quantification of wild-type full-length RFP INF2 localization pattern in podocytes. PER, Peri-nuclear ER-like pattern; MR, membrane-rich pattern; DC, diffused cytoplasmic pattern. Color bars correspond to the coexpression of GFP-and RFP-tagged INF2 conditions. Red bars, coexpression with Full-length GFP-INF2; Blue bars, coexpression with N-fragment GFP-INF2; Green bars, coexpression with a noncleavable GFP-INF2. Inlet grey bars represent their respective controls (without RFP-tagged INF2 expression). The peri-nuclear ER-like and membrane-rich localization pattern of wild-type RFP-INF2 is altered to a diffused cytosolic pattern of localization by the R218Q GFP-INF2 presence. (*p<0.01; Student’s *t-*test). (C) Representative cell images for wild-type full-length RFP-INF2 localization with various GFP-INF2 forms. Scale bar, 10 µm (D) Interaction analysis of Wild-type INF2 with R218Q INF2. Coimmunoprecipitation of wild-type HA-INF-FLAG with different GFP-tagged INF2 expression forms. GFP-tagged and HA-tagged INF2 forms were co-transfected in 293T cells. Cell lysates were pulldown using the FLAG tag and blotted for HA. HA immunoblot confirmed the immunoprecipitation of wild-type full-length INF2. GFP immunoblot showed an interaction of wild-type full-length INF2 with both wild-type and R218Q mutant forms of N-fragment, full-length and non-cleavable INF2. Each immunoblot is representative of three independent experiments with similar results. IP, Immunoprecipitation; IB, Immunoblot (C&D) Localization analysis of wild-type INF2 in the presence of R218Q INF2. (E) Pyrene actin polymerization assays containing 2 μM actin monomer (10% pyrene label), 6 μM profilin and 50 nM capping protein. Actin polymerizing activity of 20nM full-length wild-type INF2 was assessed with 0.5 μM wild-type and R218Q DID regions respectively. Neither the wild-type nor the R218Q-DID region had any effect in the actin polymerizing activity of full-length INF2.

To determine how INF2-DID mutations lead to this aberrant actin distribution, we examined the localization (Figure 2B,C) and activity of mutant INF2 in podocytes (Figure 2D,E). In podocytes, INF2-DID protein is present in a full-length INF2-CAAX isoform, and in an INF2 N-terminal cleavage product fragment ^24^. Therefore, we examined the ability of these forms of mutant INF2-DID to localize and interact with wild-type full-length INF2-CAAX in podocytes. We transiently overexpressed RFP-tagged (wild type) full-length INF2-CAAX, GFP-tagged (wild type and R218Q mutant) N-terminal fragment, full-length INF2-CAAX, and non-cleavable full-length INF2-CAAX in mouse INF2 KO podocytes. When GFP-tagged wild-type INF2 is present, the RFP-tagged wild-type full-length INF2-CAAX isoform was localized mainly at the ER-rich cell-body (PER) and some cell boundary regions (MR). In contrast, when GFP-tagged R218Q mutant INF2 isoforms are present, the RFP-tagged wild-type INF2 shifts to a diffuse cytoplasmic localization pattern (DC) (Figure 2B-C). We further probed for their interaction to confirm that these localization patterns are indeed due to the interaction between the wild-type and mutant forms of INF2 proteins. We co-expressed wild-type HA-INF2-FLAG with GFP-tagged wild-type or mutant INF2 forms in 293T cells and examined their interactions through Co-IP assays. We found that the N-fragment, full-length, and non-cleavable forms of R218Q INF2 all interact with full-length wild-type INF2-CAAX, as is the case with the equivalent wild-type forms (Figure 2D).

We also tested whether this interaction was due to a DID-DAD interaction or dimerization between N-fragment and full-length INF2. To determine that, we examined the biochemical activity of full-length wild-type INF2 with the wild-type and R218Q mutant N-terminus region. Our results showed that neither the wild-type nor the mutant form of DID regions affected the biochemical activities of wild-type full-length INF2 (Figure 2E), suggesting that the interaction is due to dimerization and the DAD activity of the wild-type full-length INF2 can remain functional post interaction. Taken together, these results show that both the N-fragment and full-length forms of INF2 are able to interact with each other and that this interaction is not noticeably altered by disease-causing mutations. However, the presence of a pathogenic mutation dramatically alters the localization of the wild-type INF2-CAAX, leading to INF2 actin polymerizing activity at abnormal sites.

### Cellular processes altered in the course of INF2-mediated FSGS

The gain-of-function effect of mutant INF2 underscores the sensitivity of podocytes to the development of functional alterations, leading to their structural and functional collapse. Therefore, we performed glomerular RNAseq transcriptome-based gene set enrichment analysis of PAN-stressed heterozygous knock-in and heterozygous knock-out mouse models to identify the altered downstream processes. We conducted our study at an early post-injury time (day 3 post PAN stress) to better focus on identifying direct changes due to mutant INF2 activity rather than more generalized effects reflecting advanced stages of injury. RNAseq analysis of unstressed heterozygous knock-in and heterozygous knock-out mouse glomeruli was also performed to elucidate the changes happening under the basal conditions in which podocyte morphology and glomerular function are grossly normal.

Principal component analysis of the total RNAseq data of all groups displayed distinct clustering of transcriptomic signature by genotype and phenotype (Figure 3A). The total number of differentially expressed genes reaching genome-wide statistical significance was ∼30 between heterozygous knock-in and heterozygous knock-out at the basal condition. In contrast, in the PAN-stressed condition, this number increased to ∼1500. This striking difference in the differentially expressed gene count suggests that heterozygous knock-in and heterozygous knock-out mouse models exhibit similar functional activity under basal conditions. However, with PAN stress, they differ significantly in their response, leading to an overt disease phenotype in heterozygous knock-in mice but not in heterozygous knock-out mice. Therefore, to understand the specific signatures associated with disease phenotype development, we conducted a GSEA analysis of the PAN-stressed heterozygous knock-in and heterozygous knock-out comparison. The enrichment results included multiple actin arrangement-related cellular function pathways, such as the “Cluster Of Actin-Based Cell Projections” within the 20 most enriched pathways (Figure 3C, Supplementary Table1). Integration of these pathways as a network revealed the enrichment of related cellular function pathways as clusters of shared genes. We noted at least two distinct clusters of pathways that are broadly related to (1) actin and cell membrane arrangement (adhesion) and (2) mitochondria structure and function (Figure 3D and E).

**Figure 3.**
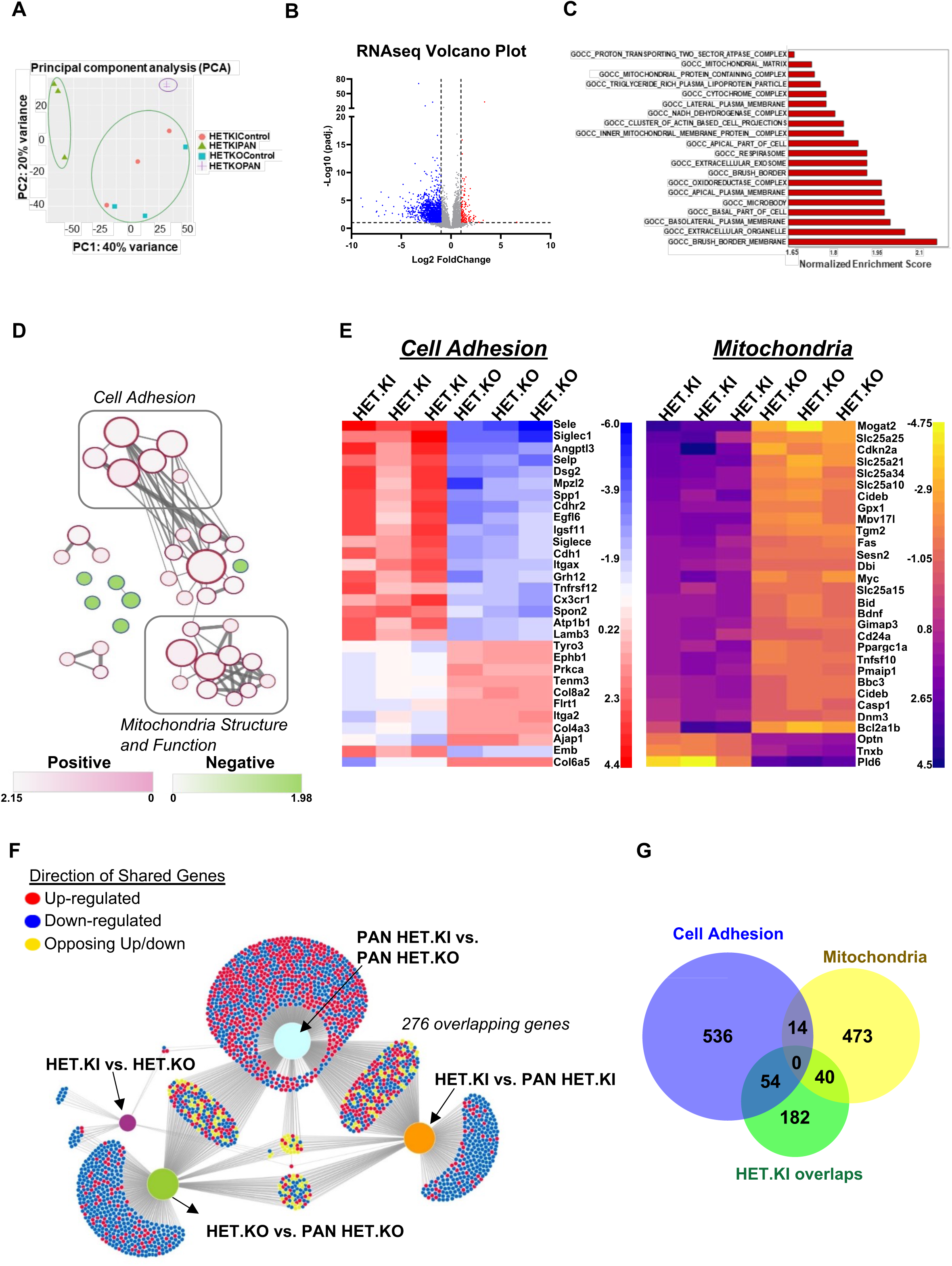
RNA-sequencing analysis identifies pathogenic processes present in INF2-related FSGS. Total RNA-seq of mouse glomeruli compared by INF2 genotype. (A) Principal component analysis plot of total RNA-seq results. (B-E) Gene set enrichment analysis of differentially expressed genes of PAN-stressed heterozygous R218Q INF2 knock-in and heterozygous INF2 knock-out mouse comparison. (B) Volcano plot of differentially expressed genes. (C) Top enrichment pathways (FDR q-value <0.05). The pathways were considered downregulated or upregulated based on the gene expression patterns of the pathway genes. (D) Enrichment network. Pathways were plotted as a network in the Cytoscape application ^44^. Color nodes indicate the upregulated and down-regulated pathways (Supplementary Table 1). Connecting lines indicate the gene overlap between pathways. Nodes were manually laid out to form a clearer picture of gene overlaps between pathways. Node clusters were identified using an Autoannotate Cytoscape application. Individual node labels were removed for clarity and to emphasize clustering. Network with individual node labeling was provided in Supplementary Figure 2. (E) The 30 most upregulated and downregulated differentially expressed genes of cell adhesion and mitochondria clusters identified throuch gene set enrichment analysis. (F) Network Visualization of differentially expressed genes (DEGs) using the Divenn application ^45^. DEGs from different comparison sets were mapped as a network to identify overlapping genes between them. A total of 276 genes were common between comparison sets that involved disease development (PAN-HET KI). (G) Venn diagram showing consensus between two methods of analysis: (i) gene enrichment analysis between PAN-stressed heterozygous knock-in versus heterozygous knock-out and (ii) Differentially expressed genes shared between comparison sets involving disease phenotype: HET-KI vs PAN-HET KI, PAN-HET KO vs PAN-HET KI.

To further confirm that these changes are robust and meaningfully associated with disease, we wished to determine if a similar conclusion is obtained with an alternate approach to RNAseq analysis. To perform this analysis, we also included the differentially expressed genes from two other comparisons: (i) heterozygous knock-in versus PAN-stressed heterozygous knock-in and (ii) heterozygous knock-out versus PAN-stressed heterozygous knock-out, and collectively analyzed all the four comparison groups to identify the overlapping gene expression changes common to these different comparisons. We reasoned that the overlapping genes between the comparison groups involving the disease phenotype are likely significant in the disease process, while others may not be relevant to disease pathology. We noted 276 overlapping genes between the comparison groups involving the PAN-stressed heterozygous knock-in mouse model (Figure 3F). Of these, 94 genes (one-third) overlap with the cell adhesion and mitochondria-related pathways identified from the gene set enrichment analysis (Figure 3G). Together, this observation suggests that these processes could be compromised due to INF2 mutation and likely play an important role in disease progression.

### Mutant mouse podocytes show altered cell adhesion and altered mitochondrial morphology

We next evaluated whether the alterations in cell adhesion and mitochondria-related processes identified through RNAseq analysis are observed in the podocytes of these mice. Our previous studies have shown that both the siRNA-silenced INF2 human podocytes and mutant podocytes from R218Q knock-in mice exhibit impaired cell spreading ^24, 29^. More recently, we have shown that N-fragment (DID region) mediates the cell-spreading function of INF2, and that this spreading may be altered in the presence of a mutation ^24^. Similarly, INF2-mediated actin arrangements have been implicated in mitochondrial fission ^30^. While these studies suggest a role for INF2 in cell spreading and mitochondria, how they compare in basal versus stressed (or disease) conditions and how disease mutations alter these processes has not been defined.

To examine this, we first tested the cell adhesion properties of mouse podocytes of various INF2 genotypes in both basal and PAN-stressed conditions using cortactin, paxillin, and F-actin staining. We conducted this analysis on a micropatterned substrate in order to have uniformity in cell dimensions and shape (Figure 4A). Under basal conditions, wild-type podocyte morphology on crossbow-shaped micropatterns showed cortactin present in lamellipodium and paxillin present in focal adhesions, with F-actin emanating from them. This distribution pattern is retained in heterozygous and homozygous INF2 knockout podocytes but was altered in both heterozygous knock-in and homozygous knock-in podocytes. We noted a loss of both lamellipodium-cortactin and focal adhesion-paxillin in homozygous knock-in podocytes, whereas in heterozygous knock-in podocytes, this is limited to the loss of cortactin in lamellipodium. We also examined how these trends manifest in response to PAN stress. We noted that the wild-type, heterozygous knock-out and homozygous knock-out podocytes retained their characteristic lamellipodium-cortactin, focal-adhesion paxillin, and F-actin distribution pattern. By contrast to the basal condition, heterozygous knock-in podocytes in the PAN-stressed condition were similar to homozygous knock-in podocytes but with more pronounced effect with loss of both lamellipodium-cortactin and focal adhesion-paxillin (Figure 4B).

**Figure 4.**
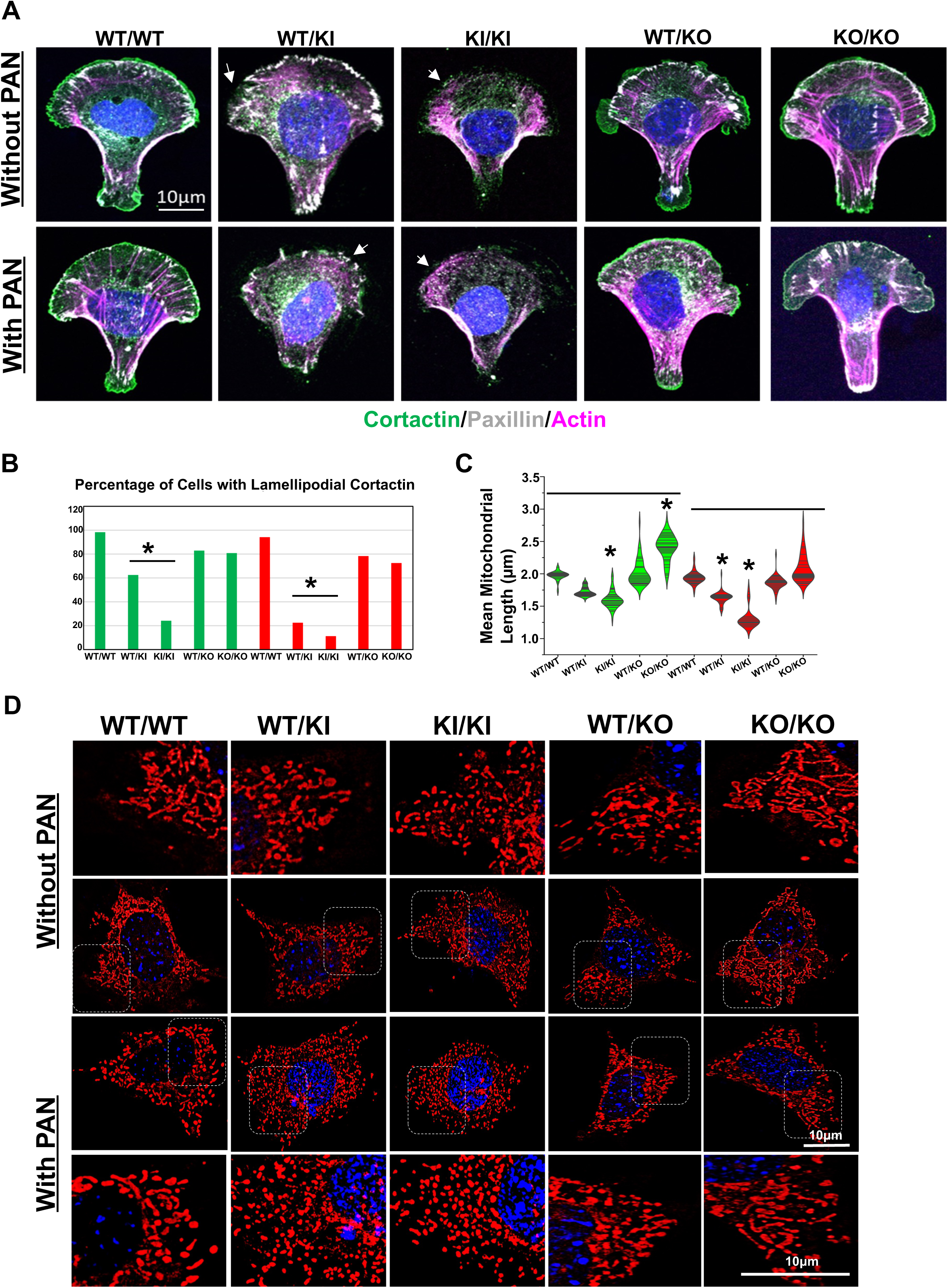
Cell adhesion and mitochondria characterization in normal and mutant mouse podocytes. (A-B) Cell adhesion assays in cross-bow micropatterns. (A) Mouse podocytes from different INF2 genotypes were compared for their cell adhesion qualities in both basal and PAN-stressed conditions using cortactin, paxillin, and F-actin staining (n, 100 cells for each group). Lack of lamellipodium cortactin, focal adhesion paxillin, and diffused cytoplasmic F-actin were associated with defective cell spreading and adhesion. R218Q knock-in podocytes showed defective cell adhesion qualities in basal and PAN-stressed conditions (white arrow). Scale bar, 10 µm. (B) Quantification for defective cell adhesion (as determined by lamellipodium cortactin). Green bars: Untreated, Red bars: PAN-treated. Heterozygous and Homozygous R218Q knock-in podocytes showed a significant defect in cell adhesion (*p>0.001 R218Q INF2 knock-in podocytes versus other groups; one-way ANOVA and Tukey’s multiple comparison test; n=100 for each group) (C-D) Mitochondria assessment using mitotracker staining. Mouse podocytes from all INF2 genotypes were compared for mitochondrial length (n, 100 cells for each group). Live cells were imaged as z-stacks and maximum-intensity projection images were generated to visualize the entire mitochondrial network in cells. (C) Quantification of mean mitochondrial length. Green: Untreated; Red: PAN-treated. Under basal conditions, R218Q INF2 homozygous knock-in podocytes had shorter mitochondria, whereas homozygous knock-out podocytes had longer mitochondria. (*p>0.001, R218Q INF2 homozygous knock-in podocytes versus other groups, homozygous knock-out podocytes versus other groups; one-way ANOVA and Tukey’s multiple comparison test; n=100 for each group) In PAN condition, heterozygous and homozygous R218Q INF2 knock-in podocytes showed shorter mitochondria. (*p>0.001, R218Q INF2 heterozygous and homozygous knock-in podocytes versus other groups; one-way ANOVA and Tukey’s multiple comparison test; n.100 for each group) (D) Representative mitochondria images from all genotypes in basal and PAN-treated conditions. Higher magnification images of highlighted regions were shown in a separate panel. Scale bar, 10 µm.

To examine mitochondria in podocytes, we compared the mitochondria filament length in basal and PAN-stressed conditions. We observed differences as a function of INF2 genotype (Figure 4C). Under basal conditions, homozygous knock-in podocytes showed shorter mitochondrial filaments than wild-type podocytes. Interestingly, homozygous knockout podocytes had longer mitochondrial filaments. The PAN stress further enhanced the trends in knock-in podocytes, making the mitochondria filaments in both heterozygous knock-in and homozygous knock-in podocytes significantly shorter than the PAN-stressed wild-type podocytes. In contrast, despite the PAN stress, heterozygous and homozygous knockout podocytes exhibited mitochondrial filament length similar to wild-type podocytes (Figure 4C and D). Together, these observations confirm that the presence of mutant alleles in podocytes leads to cell adhesion and mitochondria defects, making them vulnerable to developing disease with PAN stress.

### INF2-patient kidney organoid-derived podocytes recapitulate mouse disease phenotypes

To examine whether our results correlate with human conditions, we prepared IPSC cells from an individual with INF2-related FSGS. Sequence analysis confirmed that the IPSCs retained the heterozygous S186P INF2 mutation. We then used these cells to prepare kidney organoids and assessed their morphology using marker proteins. The results showed that the S186P INF2 IPSCs formed kidney organoids with a typical distribution of glomeruli (20%) and tubular structures (70%) determined by their respective marker proteins, nephrin, and E-cadherin. However, when we examined the sub-cellular localization of these proteins, we noted that the podocytes of wild-type organoids displayed a basolateral localization for nephrin, whereas in S186P INF2 organoids, nephrin had an altered punctate pattern mislocalizing to peri-nuclear regions. The PAN injury further altered these patterns, exhibiting a punctate cytoplasmic distribution in normal organoids and a loss of nephrin staining intensity in SI86P organoids. In contrast, the localization of E-cadherin in tubules remained similar between wild-type and S186P organoids, both in basal and PAN-treated conditions (Figure 5B).

**Figure 5.**
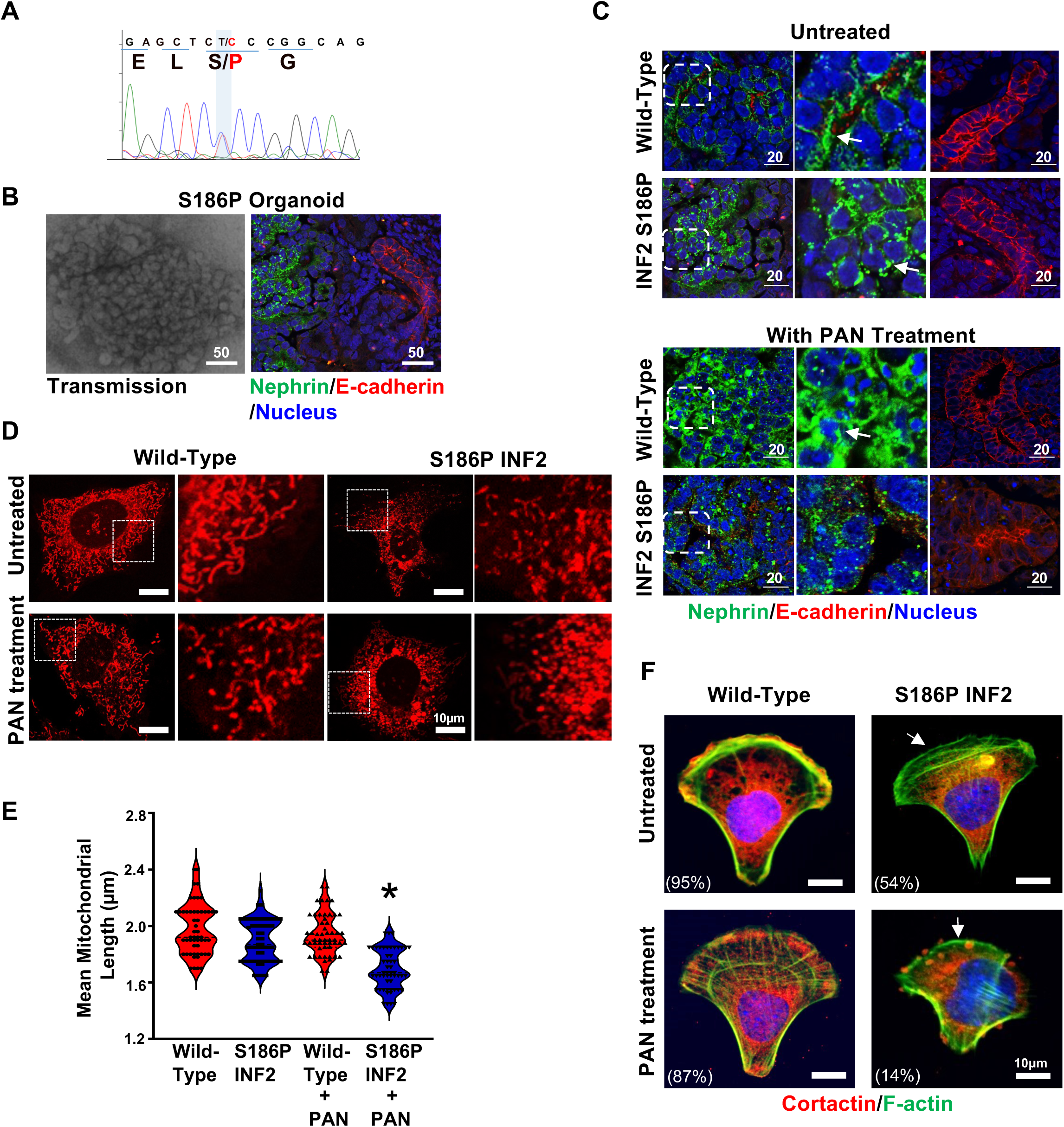
FSGS patient’s IPSC-kidney organoid-derived podocytes exhibit defective cell adhesion and mitochondria. (A) FSGS patient’s IPSC characterization. DNA sequencing analysis of the INF2-DID region showed a heterozygous missense mutation causing serine (S) to proline (P) at amino acid position 186 in INF2-CAAX. (B) S186P INF2 IPSCs formed kidney organoids with glomeruli and tubular structures. There is no apparent difference in the distribution of glomeruli (nephrin-stained) and tubules (E-cadherin-stained). (C) Marker protein analysis of S186P organoids in basal and PAN-treated condition. The basolateral localization pattern of nephrin is altered to a punctate pattern by the S186P mutation. PAN injury affected nephrin staining pattern in normal and S186P organoids. The E-cadherin staining pattern remained unaffected in tubular structures. (D-E) Mitochondria assessments. Outgrown podocytes were examined for mitochondrial filament length. PAN-treated S186P podocytes showed a shorter filament length. (*p>0.001, PAN-treated heterozygous knock-in S186P INF2 podocytes versus other groups; one-way ANOVA and Tukey’s multiple comparison test; n.100 for each group) (F) Cell Adhesion Assessments. Heterozygous S186P knock-in podocytes lack lamellipodial cortactin (white arrow) and are defective in cell adhesion on a cross-bow micropattern in basal and PAN-treated conditions. The percentage of cells with lamellipodial cortactin was indicated. (*p>0.001, Percentage of heterozygous knock-in S186P INF2 podocytes versus other groups; one-way ANOVA and Tukey’s multiple comparison test; n.100 for each group) Scale bar, 10 µm.

We also examined whether the alterations in cell adhesion and mitochondrial processes noted in mutant mouse podocytes are present in human S186P INF2 podocytes. To evaluate this, we used podocytes outgrown from glomeruli of normal and S186P organoids and assessed their cell adhesion and mitochondrial qualities. We found that wild-type podocytes were able to establish cortactin in the lamellipodium and spread normally in the cross-bow micropattern in both basal and PAN-treated conditions, whereas S186P INF2 podocytes were defective in both conditions (Figure 5C). Similarly, we found that the average mitochondrial filament length in S186P podocytes was shorter than the average mitochondria filament length in wild-type podocytes in both basal and PAN-stressed conditions. (Figure 5D). These results confirm that defects in cell adhesion and mitochondria are consistent cellular features of INF2-related FSGS.

## DISCUSSION

In this study, by comparing the injury responses of INF2 knock-in and knock-out mouse models, we have demonstrated that kidney disease occurs as a result of mutant INF2’s gain-of-function effects. Given that all of the clearly pathological INF2 mutations that have been identified to date are missense variants, this observation explains the disease’s autosomal dominant inheritance pattern. In addition, we observed that the mutant INF2 acquires gain-of-function by altering the localization of wild-type INF2, thereby leading to INF2 activity at abnormal sites. These results are consistent with earlier findings on wild-type INF2 and mutant INF2 localization patterns ^24, 31, 32^. In this model, INF2 dimers composed of wild-type and mutant INF2 will differ in their localization from wild-type INF2 dimers and induce abnormal actin cytoskeletal arrangements. By doing so, mutant INF2 can cause defects in various processes integral to glomerular structure and function.

However, the described model, particularly in the context of gain-of-function, leaves some questions that still need to be clarified. For instance, the point mutant mouse model needs a stress injury in order to develop a disease phenotype. At endogenous levels, we have noted that mutant protein appears unstable, reducing levels in the knock-in podocytes. These observations are intriguing in that it would be reasonable to hypothesize that INF2 is a stress response protein, requiring a stress-induced injury for its function to become essential. Under such a model, we would expect reduced or absent INF2 to cause disease due to loss of some necessary INF2 function. However, to the contrary, the INF2 knock-out mouse model does not develop kidney disease, arguing against a loss-of-function mechanism. However, if the human (and mouse) INF2-associated disease is driven solely by gain-of-function, the evolutionary pressure (physiological need) to maintain high INF2 expression in podocytes remains unclear.

The observation that disease-causing mutations are all located in the N-terminal region requires explanation. Previously we showed that INF2 is cleaved, and post-cleavage, the INF2 N-terminus region can regulate the DAD activity of mDIA formins ^24^. While this provides one possible mechanism in which the INF2 N-terminal region may function independently of the C-terminus, other regulatory mechanisms still need to be described. In this study, our analysis of INF2 localization indicates that both the N-fragment and full-length INF2, when mutated, can alter the localization of full-length wild-type INF2. This suggests that the INF2 N-terminus region has an essential function in regulating INF2 localization. Together, the results indicate at least two unique functions to the N-terminus region: (1) determining the localization of INF2-forms in cells and (ii) regulating the DAD activity. The pathogenic mutations appear to alter both of these activities.

A major goal of our RNA-seq studies was to identify the downstream events in which the mutation-induced alterations have an effect. Our findings from the gene-set enrichment analysis indicated actin and cell-membrane arrangements (grouped as cell adhesion) and mitochondria-related genes are significantly altered during the disease. These data align with the clinical observation that podocytes in many forms of progressive glomerular diseases are often defective in their adhesion with capillary tuft and manifest mitochondrial dysfunction ^33, 34^. However, we note that adhesion- and mitochondria-related processes are not the only enrichment identified and, thus, may not be the only driving factors for INF2-related disease onset and progression. We noted many other processes, related or unrelated to INF2, in our gene-enrichment analysis. For example, Madrid et al. reported that INF2 interacts with MAL2, a vesicular protein, and regulates basolateral-to-apical transcytosis in kidney epithelial cells ^35^. In our gene-enrichment analysis, we have also identified processes relating to vesicle transport being downregulated. It is thus interesting to note that other processes beyond cell adhesion- and mitochondria-cluster may also contribute to the disease progression. Future studies of such processes will establish their relation with INF2 and their significance in disease progression.

Our previous studies have led us to hypothesize that INF2 functions downstream of Rho A and inhibits mDIA-related DAD activity through the N-terminal region ^24, 29, 36, 37^. By doing so, INF2 mediates cell adhesion by promoting lamellipodial structure formation. The results shown in this study agree with this hypothesis in that the presence of mutation affects cell spreading in micropatterns. However, INF2 knock-out podocytes did not manifest similar defects in the micropatterns, suggesting that mutant INF2-driven activities are essential for the defective phenotype. Taken together, we propose that mutant podocytes may exhibit multiple alterations; in one aspect, the absence of INF2 at proper localization leads to alterations, as noted with mDIA regulation, and with the other, the presence of mutant INF2 at aberrant sites causes unwarranted alterations, which is evident in abnormal F-actin/G-actin distribution in cells (Figure 2A). Future comparative assessment of actin distribution among various sub-cellular compartments of INF2 mutant and INF2 knock-out cells will further confirm this point and help us to identify the aberrant sites where mutant INF2 may cause alterations within the cells.

Several studies have shown that INF2 and mitochondrial fission are related and how DID mutations may affect this relationship ^30, 38, 39, 40^. However, this work presents the first clear evidence for gain-of-function effects in mitochondria fission by pathogenic R218Q INF2 in podocytes. On the one hand, it is conceivable that the alteration in fission may lead to mitochondrial dysfunction and related FSGS; on the other hand, it raises the question as to why the R218Q INF2 mouse model does not develop FSGS without additional toxic injury. As discussed above, one explanation is that mutant protein clearance may limit the extent to which mitochondrial fission occurs, preventing FSGS development. It is worth noting that the knock-out podocyte manifests decreased mitochondrial fission and has altered mitochondria structure. However, our mouse data aligns with a model in which podocyte dysfunction is enhanced in the presence of the excessive fission due to point mutant R218Q rather than the decreased fission conditions due to the loss of INF2. Future studies on the podocyte mitochondrial function with excessive or reduced fission can clarify these questions.

We have found that S186P INF2 kidney organoids, derived from the INF2 patient’s IPSC, exhibit an altered nephrin distribution from wild-type kidney organoids. Yoshimura et al. and others have shown that nephrin exhibits a basolateral distribution in normal organoids, and this pattern may be altered as a consequence of mutations in various genes ^41, 42^. Meanwhile, in mouse models, nephrin distribution remains unaltered in basal conditions in both R218Q INF2 knock-in and INF2 knock-out models ^24, 36^. However, R218Q INF2 becomes defective when recycling nephrin to the plasma membrane when injured ^43^. Reconciling these disparate results will require assessments from a matured vascularized organoid system with in vivo physiology. Our findings suggest that the effect of a S186P INF2 mutation is similar to the effect of R218Q in causing impaired cell adhesion and mitochondria structure.

In summary, we have shown that mutant INF2’s gain-of-function activity drives INF2-related FSGS (Figure 6). The mutant INF2 acquires this quality by altering the localization of wild-type INF2, thereby making INF2 exert activity at aberrant sites. In addition, we have observed that cell adhesion and mitochondrial structures are significantly affected by pathogenic mutations in INF2. These data provide a plausible explanation for the autosomal dominant inheritance pattern of the disease and highlight the molecular events driving the INF2-related FSGS. Future studies tailored to define the full range of INF2-related pathways and their significance in maintaining the podocyte’s unique structure will increase our understanding of INF2 in normal physiology and disease pathology.

**Figure 6.**
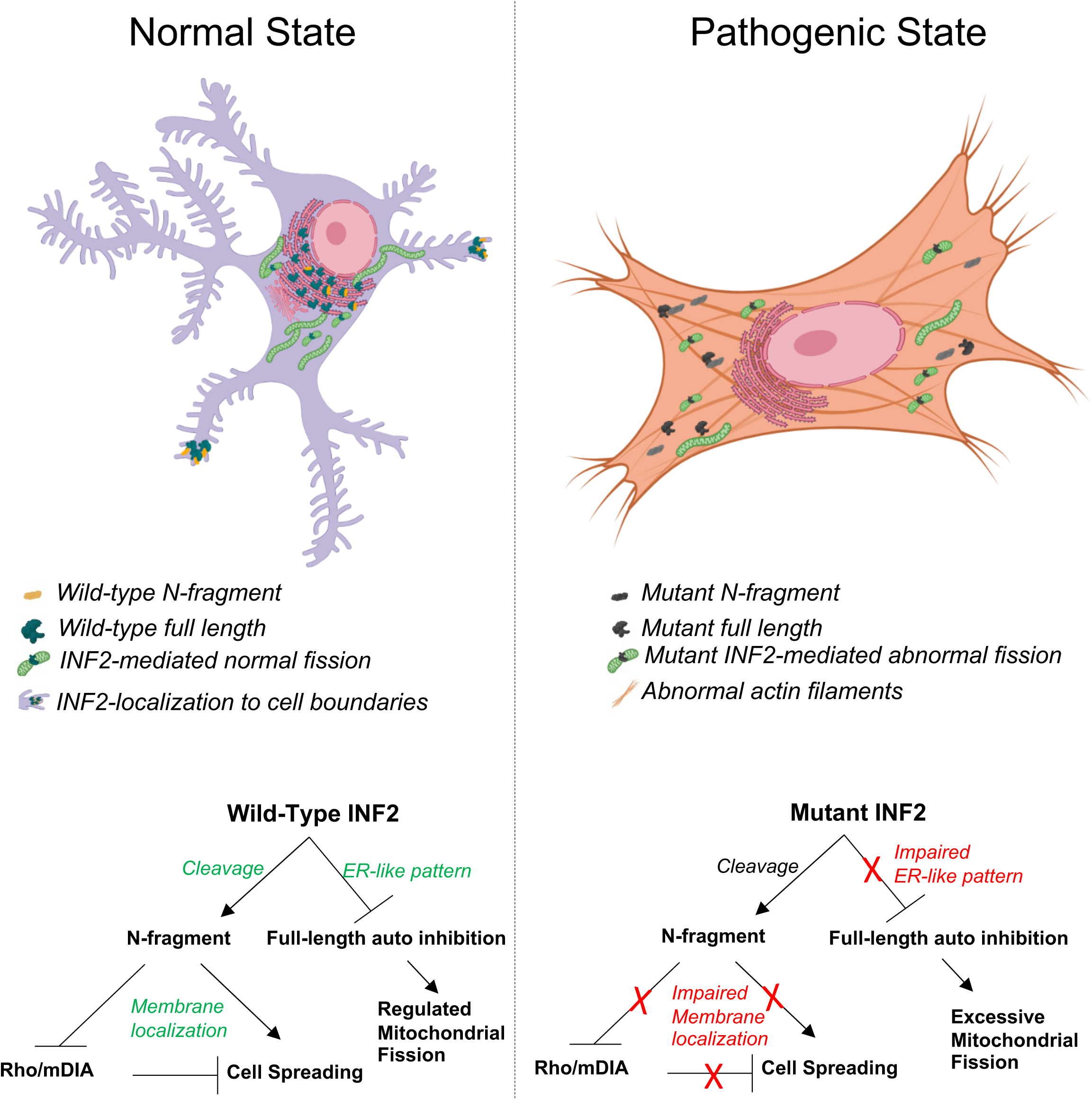
Summary model for INF2-related FSGS. INF2-DID region exists in two INF2 forms: Full-length INF2 and N-fragment INF2. Full-length INF2 predominantly localizes to peri-nuclear ER-rich regions, while N-fragment preferentially localizes to membrane regions. However, with the pathogenic mutations, their localization is altered to a diffused cytoplasmic pattern. In one possible model for disease development, the pathogenic mutation may cause (1) abnormal actin distribution due to INF2’s activity at altered localization sites and (2) excessive mitochondria fission. With an extraneous insult that causes podocytes to lose their structural integrity, the N-fragment may not localize properly, perhaps unable to counteract mDIA signaling, causing defective adhesion and impaired recovery. This combination of mutant-driven predisposition and post-injury defect may lead to chronic pathology and FSGS.

## ACKNOWLEDGEMENTS

We thank BIDMC’s confocal imaging and histology facility for technical assistance with experiments. We also thank Harvard stem cell core facility for IPSC preparation and characterization.

## IACUC approval

All animal experiments were approved by the Beth Israel Deaconess Center’s Institutional Animal Care and Use Committee (IACUC). The studies were performed in accordance with institutional and national guidelines for Animal Care and Use. The study is compliant with all relevant ethical regulations regarding animal research.

## IRB approval

The study was approved by the Institutional Review Board (IRB) of Beth Israel Deaconess Medical Center (Protocol No.# 2009P000430). Blood samples were collected and handled according to the guidelines described in the Code of Conduct for Proper Use of Human Tissue, with written informed consent from the patients prior to sampling.

## DATA SHARING

Original data files have been uploaded in SRA (PRJNA1119066). Other analyzed lists of data are available from authors upon request.

## FIGURE LEGENDS

**Supplementary Figure 1.**
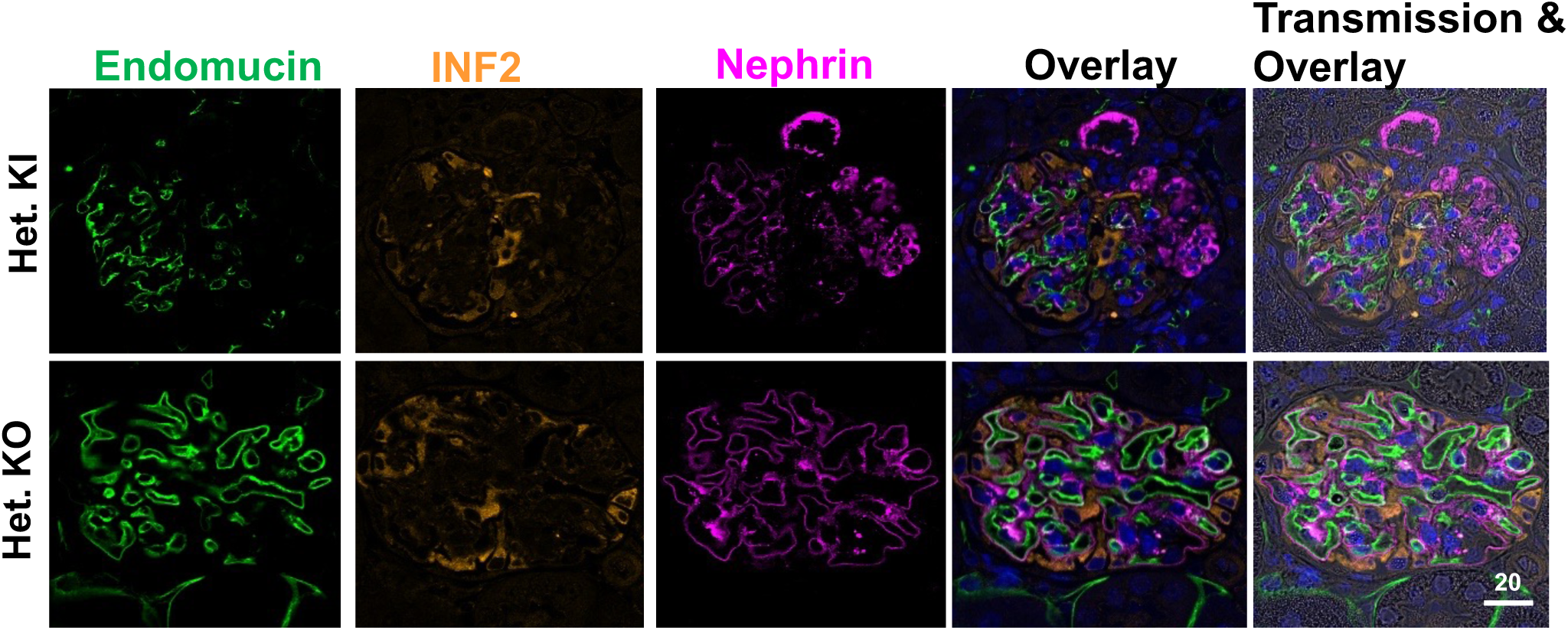
Glomerular marker protein analysis. PAN-stressed heterozygous knock-in and heterozygous knock-out mice kidney sections were stained for glomerular marker proteins: Nephrin (Purple); Endomucin (Green); INF2 (Orange). Focal sclerotic lesions were present in PAN-stressed heterozygous knock-in mice (white arrow). Scale bar, 10 µm

**Supplementary Figure 2.**
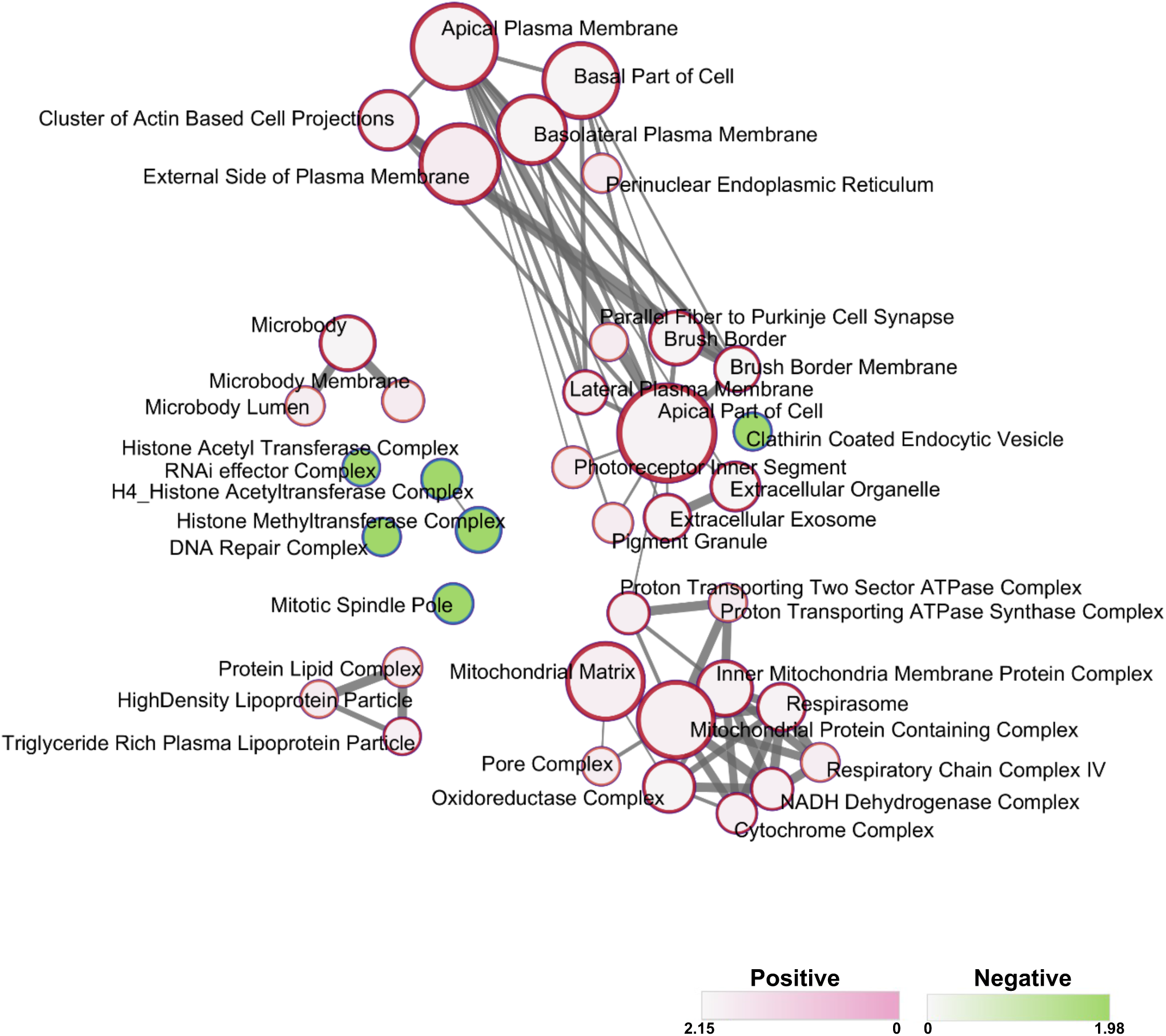
Gene Set Enrichment Network. Pathways that were significantly enriched in the comparison between the PAN-stressed heterozygous knock-in and heterozygous knock-out condition were plotted as a network in the Cytoscape application. Color nodes indicate the upregulated and down-regulated pathways. Connecting lines indicate the gene overlap between pathways. Nodes were manually laid out to form a clearer picture of gene overlaps between pathways. Individual node labels indicate the enriched pathway.

**Supplementary Table 1**. **List of enrichment pathways.**

