## Supplemental Table 1 for "Missense Mutant Gain-of-Function Causes Inverted Formin 2 (INF2)-Related Focal Segmental Glomerulosclerosis (FSGS)"

**Supplementary Table 1. Gene sets enriched in Het-KI phenotype**

| Gene Sets | Size | NES | FDR q-value |
|---|---|---|---|
| GOCC_BRUSH_BORDER_MEMBRANE | 72 | 2.16 | 0 |
| GOCC_EXTRACELLULAR_ORGANELLE | 96 | 2.05 | 0 |
| GOCC_BASOLATERAL_PLASMA_MEMBRANE | 229 | 2 | 0.001 |
| GOCC_BASAL_PART_OF_CELL | 274 | 1.98 | 0.001 |
| GOCC_MICROBODY | 136 | 1.98 | 0.001 |
| GOCC_APICAL_PLASMA_MEMBRANE | 341 | 1.97 | 0.001 |
| GOCC_OXIDOREDUCTASE_COMPLEX | 112 | 1.97 | 0.001 |
| GOCC_BRUSH_BORDER | 128 | 1.92 | 0.001 |
| GOCC_EXTRACELLULAR_EXOSOME | 80 | 1.92 | 0.001 |
| GOCC_RESPIRASOME | 88 | 1.92 | 0.001 |
| GOCC_APICAL_PART_OF_CELL | 421 | 1.89 | 0.002 |
| GOCC_INNER_MITOCHONDRIAL_MEMBRANE_PROTEIN_ COMPLEX | 139 | 1.84 | 0.004 |
| GOCC_CLUSTER_OF_ACTIN_BASED_CELL_PROJECTIONS | 166 | 1.84 | 0.004 |
| GOCC_NADH_DEHYDROGENASE_COMPLEX | 47 | 1.81 | 0.006 |
| GOCC_LATERAL_PLASMA_MEMBRANE | 60 | 1.78 | 0.009 |
| GOCC_CYTOCHROME_COMPLEX | 35 | 1.78 | 0.009 |
| GOCC_TRIGLYCERIDE_RICH_PLASMA_LIPOPROTEIN_PARTICLE | 15 | 1.76 | 0.011 |
| GOCC_MITOCHONDRIAL_PROTEIN_CONTAINING_COMPLEX | 283 | 1.74 | 0.013 |
| GOCC_MITOCHONDRIAL_MATRIX | 285 | 1.73 | 0.015 |
| GOCC_PROTON_TRANSPORTING_TWO_SECTOR_ATPASE_ COMPLEX | 47 | 1.67 | 0.028 |
| GOCC_RESPIRATORY_CHAIN_COMPLEX_IV | 23 | 1.66 | 0.034 |
| GOCC_PORE_COMPLEX | 20 | 1.65 | 0.035 |
| GOCC_MICROBODY_LUMEN | 27 | 1.65 | 0.033 |
| GOCC_HIGH_DENSITY_LIPOPROTEIN_PARTICLE | 16 | 1.62 | 0.045 |
| GOCC_PROTEIN_LIPID_COMPLEX | 27 | 1.61 | 0.049 |
| GOCC_PIGMENT_GRANULE | 32 | 1.61 | 0.051 |
| GOCC_MICROBODY_MEMBRANE | 45 | 1.56 | 0.08 |
| GOCC_PERINUCLEAR_ENDOPLASMIC_RETICULUM | 26 | 1.56 | 0.081 |
| GOCC_PROTON_TRANSPORTING_ATP_SYNTHASE_COMPLEX | 20 | 1.55 | 0.081 |
| GOCC_EXTERNAL_SIDE_OF_PLASMA_MEMBRANE | 303 | 1.55 | 0.08 |
| GOCC_PARALLEL_FIBER_TO_PURKINJE_CELL_SYNAPSE | 17 | 1.54 | 0.086 |
| GOCC_PHOTORECEPTOR_INNER_SEGMENT | 42 | 1.53 | 0.091 |
| GOCC_RIBOSOMAL_SUBUNIT | 193 | 1.49 | 0.134 |
| GOCC_MICROVILLUS | 85 | 1.49 | 0.131 |
| GOCC_CELL_PROJECTION_MEMBRANE | 260 | 1.49 | 0.129 |
| GOCC_TRICARBOXYLIC_ACID_CYCLE_ENZYME_COMPLEX | 20 | 1.47 | 0.149 |
| GOCC_APICOLATERAL_PLASMA_MEMBRANE | 22 | 1.47 | 0.146 |
| GOCC_ENDOPLASMIC_RETICULUM_GOLGI_INTERMEDIATE_ COMPARTMENT_MEMBRANE | 16 | 1.46 | 0.161 |

| | | | |
|---|---|---|---|
| GOCC_CYTOSOLIC_LARGE_RIBOSOMAL_SUBUNIT | 62 | 1.44 | 0.173 |
| GOCC_APICAL_JUNCTION_COMPLEX | 129 | 1.44 | 0.174 |
| GOCC_PROTON_TRANSPORTING_TWO_SECTOR_ATPASE_ COMPLEX_CATALYTIC_DOMAIN | 16 | 1.44 | 0.171 |
| GOCC_VACUOLAR_PROTON_TRANSPORTING_V_TYPE_ATPASE _COMPLEX | 23 | 1.43 | 0.173 |
| GOCC_CATENIN_COMPLEX | 16 | 1.43 | 0.17 |
| GOCC_MONOATOMIC_ION_CHANNEL_COMPLEX | 153 | 1.43 | 0.169 |
| GOCC_LARGE_RIBOSOMAL_SUBUNIT | 119 | 1.43 | 0.167 |
| GOCC_CYTOSOLIC_RIBOSOME | 123 | 1.42 | 0.183 |
| GOCC_PROTON_TRANSPORTING_V_TYPE_ATPASE_COMPLEX | 27 | 1.41 | 0.186 |
| GOCC_TRANSPORTER_COMPLEX | 258 | 1.41 | 0.183 |
| GOCC_HIPPOCAMPAL_MOSSY_FIBER_TO_CA3_SYNAPSE | 42 | 1.41 | 0.182 |
| GOCC_SMALL_RIBOSOMAL_SUBUNIT | 79 | 1.4 | 0.193 |
| GOCC_CYTOSOLIC_SMALL_RIBOSOMAL_SUBUNIT | 44 | 1.39 | 0.205 |
| GOCC_TIGHT_JUNCTION | 114 | 1.39 | 0.208 |
| GOCC_PHOTORECEPTOR_OUTER_SEGMENT | 52 | 1.38 | 0.214 |
| GOCC_CATION_CHANNEL_COMPLEX | 100 | 1.36 | 0.252 |
| GOCC_PROTON_TRANSPORTING_TWO_SECTOR_ATPASE_ COMPLEX_PROTON_TRANSPORTING_DOMAIN | 21 | 1.36 | 0.25 |
